# Tackling Mitosis Domain Generalization in Histopathology Images with Color Normalization

**DOI:** 10.1101/2022.08.23.505051

**Authors:** Satoshi Kondo, Satoshi Kasai, Kousuke Hirasawa

**Affiliations:** Muroran Institute of Technology, Hokkaido, Japan; Niigata University of Healthcare and Welfare, Niigata, Japan; Konica Minolta, Inc., Osaka, Japan

**Keywords:** Mitosis detection, Domain adaptation, Color normalization

## Abstract

In this report, we propose a method for mitosis detection in histopathology images of MIDOG2022 challenge dataset. Our method is based on unsupervised domain adaptation at input-level, and a combination of color normalization and object detection. We evaluate our method by using the preliminary test set including 20 cases and obtain 0.716 F1 score.

## Introduction

Mitosis detection is a key component of tumor prognostication for various tumors. In recent years, deep learning based methods have shown high detection accuracy for mitosis on the level of human experts (1). Mitosis is known to be relevant for many tumor types, however, when a model is trained on a specific tumor or tissue type, the performance of the model typically drops significantly on other types.

The Mitosis Domain Generalization Challenge 2021 (MIDOG2021) (2) was held to evaluate the performance of several methods for mitosis detection by using breast cancer cases digitized by different scanners. MIDOG2022 (3) is held as extension of MIDOG2021. In MIDOG2022, histopathology images acquired from different laboratories, from different species (human, dog, cat) and with five different whole slide scanners are newly provided. The training set consists of 400 cases and contains at least 50 cases each of prostate carcinoma, lymphoma, lung carcinoma, melanoma, breast cancer, and mast cell tumor. The test set contains 10 cases each of 10 tumor types.

Our proposed method is a kind of unsupervised domain adaptation (UDA). UDA is a type of domain adaptation and exploits labeled data from the source domain and unlabeled data from the target one (4). UDA approaches can be grouped, based on where the domain adaptation is performed, at input-level, at feature-level, at output-level or at some ad-hoc network levels (5). Our proposed method performs the domain adaptation at input-level.

## Proposed Method

Figure reffig:overview shows the overview of our proposed method. Our method is two-step approach. The first step is color normalization as domain adaptation. The color normalized image is divided into patches and object detection is performed for each patch. The detection results for the patches are combined to obtain the final detection results.

We employ the method proposed in (6) for color normalization. In (6), an input image is first decomposed into stain density maps that are sparse and non-negative. This is based on an assumption that tissue samples are stained with only a few stains and most tissue regions are characterized by at most one effective stain. For a given image, its stain density maps are combined with stain color basis of a target image, thus altering only its color while preserving its structure described by the maps.

Input histopathology images are whole slide images and these sizes are very large. Therefore, the images after the color normalization are divided into patches since it is hard to process a whole slide image at once. We use an object detector to perform mitosis detection for each patch. We employ Efficientdet (7) as our object detector. The mitosis detection results for patches are merged into an original whole slide space.

## Experiments

The dataset used in our experiments includes digitize microscopy slides from 520 tumor cases, acquired from different laboratories, from different species (human, dog, cat) and with five different whole slide scanners. The training set consists of 400 cases and contains at least 50 cases each of prostate carcinoma, lymphoma, lung carcinoma, melanoma, breast cancer, and mast cell tumor. The validation dataset contains five cases each of four different tumor types.

We select one image from the training dataset as the template for the color normalization. Examples of color normalization are shown in Fig. 2. The size of a patch is 512 *×* 512 pixels and adjacent patches have overlap region in 67 pixels. In the dataset, two types of objects are annotated. One type is “mitotic” class, and an-other type is “non-mitotic (hard negative)” class. We train our object detector to detect these two types of classes. In inference, we only use mitotic class and select regions having scores higher than a threshold, 0.5. In the training of the object detector, the learning rate is 0.002, the batch size is 8, the number of epochs is 30. The optimizer is Adam (8). We use horizontal and vertical flips as augmentation. Although unlabeled images are included in the training data, we do not use those images for training. And we do not use any additional data other than dataset provided by the MIDOG2022 organizers. The evaluation is performed by using a preliminary test set. The preliminary test set uses four tumor types, each with 5 cases (20 cases in total). The evaluation is performed by using a preliminary test set. The preliminary test set uses four tumor types, each with 5 cases (20 cases in total). The experimental results show that mAP is 0.575, recall is 0.694, precision is 0.739 and F1 score is 0.716.

**Fig. 1.**
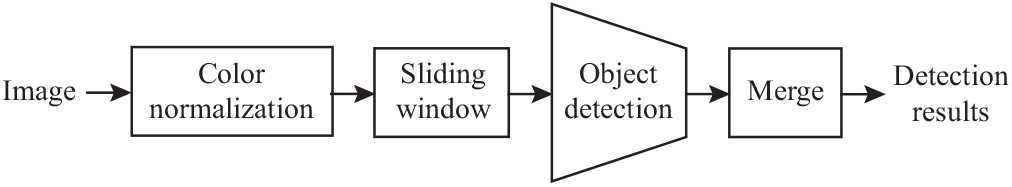
Overview of our proposed method.

**Fig. 2.**
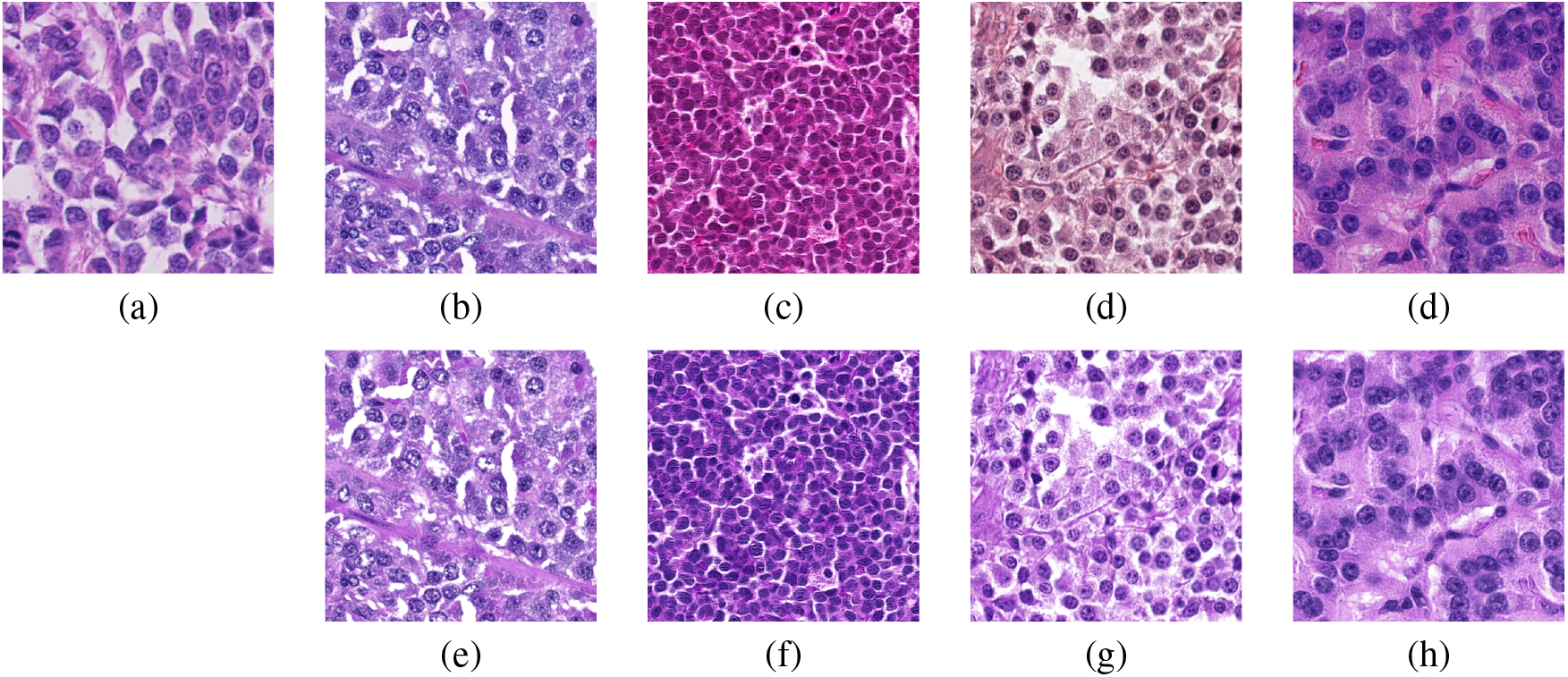
Examples of color normalization. (a) Template image for color normalization. (b)-(d) Original images. (e)-(h) Color normalized images. Each image in the bottom row corresponds to an image in the upper row and the same column.

## Conclusion

In this report, we propose a method for mitosis detection in histopathology images. Our method is based on unsupervised domain adaptation at input-level. We evaluated our method by using the preliminary test set including 20 cases and we obtained 0.716 F1 score.

